# *KAI2*-KL signalling regulates root hair elongation under magnesium deficiency by activating auxin, ethylene, and nitric oxide biosynthesis and signalling genes

**DOI:** 10.1101/2022.07.20.500783

**Authors:** Faheem Afzal Shah, Jun Ni, Xue Chen, Caiguo Tang, Lifang Wu

## Abstract

Root hair elongation (RHL), which expands the absorptive surface area of the root, is a crucial adaptation in plants for survival under magnesium (Mg) deficient soil. Despite the significance of this trait, the molecular mechanism in Mg starvation regulating RHL is elusive. We demonstrated that karrikins regulate RHL under a limited supply of Mg via crosstalk with auxin, ethylene, and NO. We used *KAI2*-KL-signalling mutants, auxin, ethylene, and NO-related genes mutants *Arabidopsis* and pharmacological method to investigate the role of *KAI2*-KL-signalling, and its interaction with ethylene, auxin, and NO in the process of RHL in Mg deficient conditions. Mg deficiency could not enhance RHL in *KAI2*-KL-signalling mutants such as *kai2* and *max2 Arabidopsis.* Interestingly, exogenous application of ethylene, nitric oxide, or auxin recovered RHL of *kai2* and *max2 Arabidopsis* under Mg deficiency. In contrast, exogenous supplementation of KAR_1_ could not rescue RHL in auxin, ethylene, and NO-related mutants *Arabidopsis*. In conclusion, we suggest that karrikins signalling might regulate the RHL in response to low Mg by acting as an upstream signalling pathway of auxin, ethylene, and NO signalling.

## Introduction

Magnesium (Mg) is a secondary nutrient that is involved in numerous fundamental biochemical processes, such as protein synthesis, chlorophyll synthesis, enzyme activation, and photosynthetic carbon fixation (Lilley and Walker, 1974; Marschner, 2005; Volker *et al*., 2005; Zheng *et al*., 1993). Thus, Mg deficiency could reduce the yield of crops (Christian *et al*., 2010). Mg starvation is a severe limitation in crop production in sandy or acidic soils because the soluble Mg^2+^ rapidly leaches from acidic or sandy soils. Mg absorption could be antagonized by potassium during uptake by plant roots (Petra and Zed, 2012; Xie *et al*., 2021). Elongation of root hairs (RH) is an adaptive mechanism in plants for survival in low nutrient’s availability, thus, low Mg availability has also been reported to have a role in RH elongation (Niu *et al*., 2014). Root hairs are the single-cell extension tubes in root epidermal cells that are dynamically modified according to their surroundings, allowing the root to perform its maximum functions in uncertain soil conditions (Brian and Helena, 2001; Emily *et al*., 2017; José *et al*., 2003). Plant hormones such as auxin, ethylene, and nitric oxide are necessary for developing RH in response to Mg availability (Ishfaq *et al*., 2021; Miao *et al*., 2017; Miao *et al*., 2018)

Ethylene is one of the key phytohormones regulating root hair elongation (Liam, 2001; Mimi *et al*., 1995). A family of genes such as 1-amino-cyclopropane-1-carboxylate synthase (ACS), which catalyze the ethylene formation, have a big influence on the root hair development. For example, *Arabidopsis* mutant in *acs7* produced shorter RH as compared to wild-type *Arabidopsis* (Carbonnel *et al*., 2020; Jason *et al*., 1998). Moreover, pharmacological studies showed that exogenous treatment of ethylene precursor1-aminocyclopropane-1-carboxylic acid (ACC) promoted the RHL, while ethylene biosynthesis inhibitors such as Ag^+^ substantially lessen RHL (Zhu *et al*., 2006). Along with ethylene, nitric oxide (NO) is an important player regulating root hair development (Chandrika *et al*., 2013). Exogenous treatment of sodium nitroprusside (SNP), “NO donor,” stimulated *Arabidopsis’s* RH initiation and elongation. In contrast, 2-(4-carboxyphenyl)-4,4,5,5-tetramethylimidazoline-1-oxyl-3-oxide (c-PTIO), which is a well-known scavenger of NO, inhibited RHL (Cristina *et al*., 2006). Besides ethylene and NO, auxin is also one of the major players in root hair elongation (Habets and Offringa, 2014; Soeno *et al*., 2010). In *Arabidopsis*, PIN-FORMED (PIN) efflux transporter such as *PIN2* and influx transporter such as *AUXIN-RESISTANT (AUX1)/LAX (*like *AUX1)* mediates the auxin transport; therefore, a mutation in *pin2* or *aux1* exhibited shorter root hairs, as compared to WT *Arabidopsis* (Křeček *et al*., 2009; Swarup and Péret, 2012).

The karrikins signalling pathway is a newly discovered plant hormone that has been found in the phenomenon of seed germination activation after a forest fire(Yao and Waters, 2020). Karrikins (KARs) are the small butenolide produced during a forest fire and activate seed germination in many plant species (Flematti *et al*., 2004). The plants, such as *Arabidopsis*, contain a karrikins protein *KARRIKIN INSENSITIVE2 (KAI2)*, which binds to *MORE AXILLARY BRANCHING2* (*MAX2*) and activate the ubiquitination of *SUPPRESSOR of MAX2 1 (SMAX1)* and *SMXL2*, consequently activates multiple downstream processes (Khosla *et al*., 2020; Lei *et al*., 2020). *SMAX1* and *SMXL2* have been reported to be involved in seed germination, cotyledons, hypocotyls, and root hair development (José Antonio *et al*., 2019; Nelson *et al*., 2012; Nelson *et al*., 2010; Stanga *et al*., 2016; Waters *et al*., 2012). Recently, a study showed that *KAI2* and *MAX2* regulate RHL in *Arabidopsis* (José Antonio *et al*., 2019). Moreover, *KAI2* signalling mediated RHL in *Arabidopsis* under phosphate starvation conditions via triggering the ethylene-*AUX1/PIN2-auxin* cascade (José Antonio *et al*., 2022).

This report revealed that Mg deficiency could not promote the RHL in *kαi2 Arabidopsis*. Exogenous treatment of KAR_1_ induced the expression of the biosynthesis, transport or signalling genes of ethylene, auxin, and nitric oxide in *Arαbidopsis* grown under Mg deficiency. We postulated that karrikins regulated root hair growth under Mg starvation by targeting multiple signalling pathways. We used ethylene biosynthesis (*acs7*) and receptor (*etr1*) genes; NO biosynthesis (*nos1* and *nia1nia2*) genes; and auxin efflux and influx carrier (*pin2, and aux1*, respectively) genes to precisely investigate the interactions of karrikins with ethylene, auxin, and NO in controlling the root hair growth. Furthermore, we exposed that Mg starvation raised the endogenous concentrations of ethylene, NO, or auxin in *Arabidopsis* roots. Our study showed that karrikins signalling might act upstream of ethylene, NO, and auxin signalling in RH elongation in *Arabidopsis* growing in Mg starvation. This study addressed the molecular mechanisms in karrikins signalling regulation of RHL under Mg starvation.

## MATERIALS AND METHODS

### Plant material and growth conditions

The *Arabidopsis* wild ecotype line (Col-0), and mutants such as *aux1* (N9583), *pin2* (N8058), *nos1* (N6511), *nia1nia2* (N6512), *acs7* (CS16570), *etr1* (N71769), *kai2 (Nelson et al., 2011), max2-2* (N9566), *smax1*(N660047), and *smxl2* (N825409), were purchased from the European *Arabidopsis* Stock Centre (http://Arabidopsis.info/). The primers were designed using NCBI Primer Blast to clone the *KAI2, MAX2, SMAX1*, and *SMXL2* (Table S**1**). Full-length sequences were double digested and were linked to the expression vector pOCA30 containing the CaMV35S promoter. The resultant overexpression vectors having a specific gene were separately transformed into the *Agrobacterium* EHA105 strain. Gene-specific overexpression lines of *Arabidopsis* were selected by gene transformation using the floral dip method. All seeds were surface sterilized by the previously used method (Shah *et al*., 2021), plated on ½ nutrients containing Petri plates, and kept at 4°C in a refrigerator for 48 hours. After two days of incubation at cold temperature, Petri plats were shifted to a growth chamber with 80% humidity, 120 mol photons m^-2^ s^-1^ light intensity, and a daily light/temperature cycle of 16 h/22 °C day and 8 h/20 °C at night. We used 4-day-old seedlings of uniform size in all treatments. The formulations of the Hoagland medium was as follows: 250 μM (NH_4_)2SO_4_, 1000 μM CaCl_2_, 500 μM MgSO_4_, 1500 μM KNO_3_, 500 μM NaH_2_PO_4_, 10 μM H_3_BO_3_, 0.5 μM ZnSO_4_, 0.5 μM MnSO_4_, 25 μM Fe-EDTA, 0.1 μM (NH_4_)_6_Mo_7_O_24_, and 0.1 μM CuSO_4_ (Hoagland and Arnon, 1938), 0.5 % (w/v) sucrose, pH of the nutrients medium was adjusted to 7 with NaOH and jelled with 0.7% (w/v) Agar (Sigma-Aldrich). In this study, we used MgSO_4_ free as Mg deficient medium (-Mg) and 500 μM MgSO_4_ (+Mg) nutrients medium (Miao *et al*., 2017). Furthermore, we added an ethylene source (Ethephon, 100 μM), ethylene activity inhibitor such as silver nitrate (AgNO_3_, 5 μM), 50 μM sodium nitroprusside (SNP, a source of nitric oxide), 100 μM cPTIO (NO scavenger), 0.1 μM Indole-3-acetic acid (IAA), 20 μM 2,3,5-triiodobenzoic acid (TIBA, auxin transport inhibitor), 1 μM Karrikin1 (KAR_1_), and 1 μM Karrikin2 (KAR_2_). We used 15 independent replicates in each treatment. Four-day-old uniform seedlings were shifted to the above-mentioned nutrient media for five days.

### Microscopy

*Arabidopsis* root hairs were measured following the previous study (Miao *et al*., 2018). Briefly, *Arabidopsis* was grown into the desired medium for five days, and RHL was calibrated between 2-3 mm from the root tip using ImageJ (Version 1.5). Micrographs were recorded using an Olympus SZX10 stereomicroscope coupled with TUCSEN 6.0 colour camera. RHL was measured using ImageJ (Version 1.5).

### Measurement of ethylene production in roots

We used the previously used method by Tian *et al*. (2009) to measure the ethylene concentration in *Arabidopsis*. In detail, 200 milligrams of roots were harvested in five ml vials having 0.7% agar medium (1 ml), and vials remained unsealed for one hour to mitigate the wounding effect. Afterwards, we sealed the vials with gas-tight stoppers. After 6 hours, the headspace gas was collected by a one ml injection and injected into a gas chromatograph (GC-7AG; Shimadzu, Tokyo, Japan) equipped with an alumina column.

### IAA determination in the roots

The quantification of the IAA was done by a previously described method by (Swarup *et al*., 2007). In detail, frozen samples of *Arabidopsis* roots were ground in liquid nitrogen using a PTFE tissue grinder, homogenized in 1.5 ml of 1mM 2,6-di-tertbutyl-p-methylphenol solved in 80% methanol, and then centrifuged at the same speed of 5000 rpm for ten minutes at 4°C. We used C18 columns to purify the supernatant. The purified product was gas dried, liquefied in 200 ml absolute methanol, and re-dried by vacuum freeze-dryer (SCIENTZ-12N|D-Ningbo Scientz Biotechnology Co., Ltd.). Finally, the dried extract was liquified in 300 ml of phosphate buffer solution (pH 7.4) to perform the enzyme-linked immunosorbent assay (ELISA). We used Multiskan FC Absorbent Microplate Reader (Thermo Fisher Scientific, Waltham, Massachusetts, U.S.) to calibrate IAA at 490 nm.

### Determination of NO levels in the roots

We used Silva *et al*. (2020) method to determine the nitric oxide. In detail, 50 mg of frozen samples of *Arabidopsis* roots were powdered in liquid nitrogen using a PTFE tissue grinder and incubated in a nitric oxide fluorescent probe called 4-amino-5-methylamino-2’,7’-difluorofluorescein (5 μM) dissolved in 50 mM phosphate buffer (pH 7.2) for one hour. Afterwards, fluorescent emission was calibrated at 515 nm emission and 495 nm excitation wavelength). 100 μM cPTIO was considered as a control.

### Gene expression Analysis

Root samples were collected from 5-week-old *Arabidopsis* shifted in Mg less or controlled medium (500 μM MgSO_4_) for 12 hours. Three samples were taken from the mixture of roots of 5 different plants of identical growth for each treatment and instantly frozen in liquid nitrogen. The 50 mg of frozen root samples were powdered by a PTFE tissue grinder, and total RNA was isolated using “A plant RNA extraction kit (Omega Bio-tek, Inc., Norcross, GA)”. RNA purification and quantification were checked through ScanDrop Nano-Volume Spectrophotometers (Analytik Jena, Jena, Germany). We used those RNA samples for qPCR, which did not smear on the agarose and have a 260/280 ratio above 2. We followed the standard protocol of “MonScript™ RTIII All-in-One Mix with dsDNase (MR00201, Monad, Wuhan, China)” and prepared the cDNA of 500 nanograms (ng) of RNA from each sample. Furthermore, these cDNA synthesized samples were further 25 times diluted with RNA-free water. RT–qPCR was prepared using the standard protocol of “QuantiNova SYBR Green PCR Master Mix (QIAGEN, Pudong, Shanghai, China)”. The qPCR was performed in “the Light Cycler®96 (Roche Diagnostics, Indiana, USA). We used the 2^-ΔΔCt^ method to measure the relative expression of the genes (Livak and Schmittgen, 2001). The statistical data were analyzed in R Studio 2022.02.0 Build 443. One-way Kruskal-Wallis test was used to investigate the effect of each treatment. Furthermore, multiple comparisons between the means were tested by post hoc Student’s t-test at P<0.05.

## Results

Magnesium starvation induces root hair elongation (RHL) by promoting the production of different hormones such as ethylene, auxin, and nitric oxide (Miao *et al*., 2018; Michael *et al*., 2009). Recently, karrikins signalling genes were discovered to be involved in RHL in *Arabidopsis* (José Antonio *et al*., 2019; José Antonio *et al*., 2022). Nevertheless, whether karrikins are involved in Mg deficiency-regulated elongation of root hairs (RH) in *Arabidopsis* remains elusive. This study discovered that the mutation in several karrikins signalling genes, or exogenously application with the pharmacological karrikins, altered the RH growth in response to Mg starvation. The results revealed that after transplantation of the 4-day-old *Arabidopsis* seedling to Mg deficient medium, *kai2* and *max2* mutants did not exhibit an increased RH length compared with wild-type plants. On the other hand, the mutants of the repressors of karrikins, such as *smax1* and *smxl2*, has significantly longer root hairs compared with *kai2, max* and wild-type *Arabidopsis* seedlings (Fig. **1a-d**). nterestingly, exogenous application of KAR_1_ and KAR_2_ significantly improved the RHL in wild type plants grown in Mg deficient medium (Fig. **1e-f**).

**Fig. 1:**
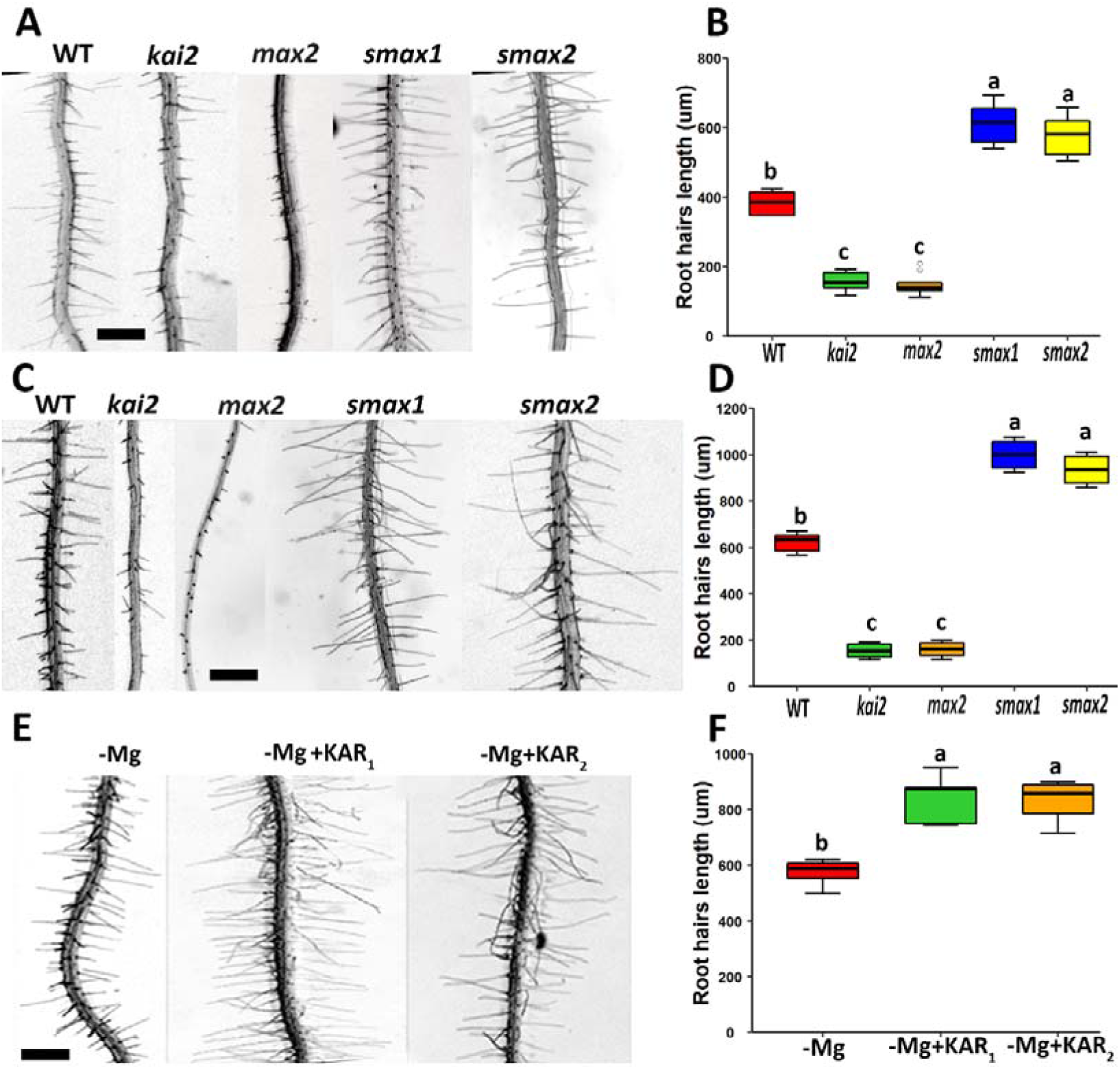
Effects of magnesium (Mg) deficiency in root hairs elongation in karrikins signalling mutants *Arabidopsis*. (**a-b**) Root hair length of karrikins signalling mutants under control condition. (**c-d**) Root hair length of karrikins signalling mutants under Mg deficient condition. (**e-f**) Impact of KAR_1_ and KAR_2_ on root hair length of *Arabidopsis* under Mg deficient condition. Four-day-old seedlings were shifted to +Mg (control) or -Mg media for five days. Each experiment was repeated three times. In (**b**), (**c**), and (**f**) Kruskal-Wallis’s test and post hoc Student’s t-test; P<0.05, n=10, were used to find the effect and significant difference between treatments; different alphabets indicate different statistical groups. Bar, 200 μm.

We then investigated the relative expression of karrikins signalling and marker genes in *Arabidopsis* grow in Mg deficient medium. The results showed that Mg starvation significantly promoted the relative expression of *KUF1, DLK2, KAI2, and MAX2* (Fig. **2a-d**). Consistently, the transcript level of karrikins functions repressing genes such as *SMAX1* and *SMXL2* was significantly decreased in *Arabidopsis* grown in the Mg deficient medium compared with Mg sufficient medium (Fig. **2e-f**). Conclusively, these results suggested that Mg status directly affected the karrikins signalling, and karrikins might be directly involved in the regulation of RHL under Mg deficiency.

**Fig. 2:**
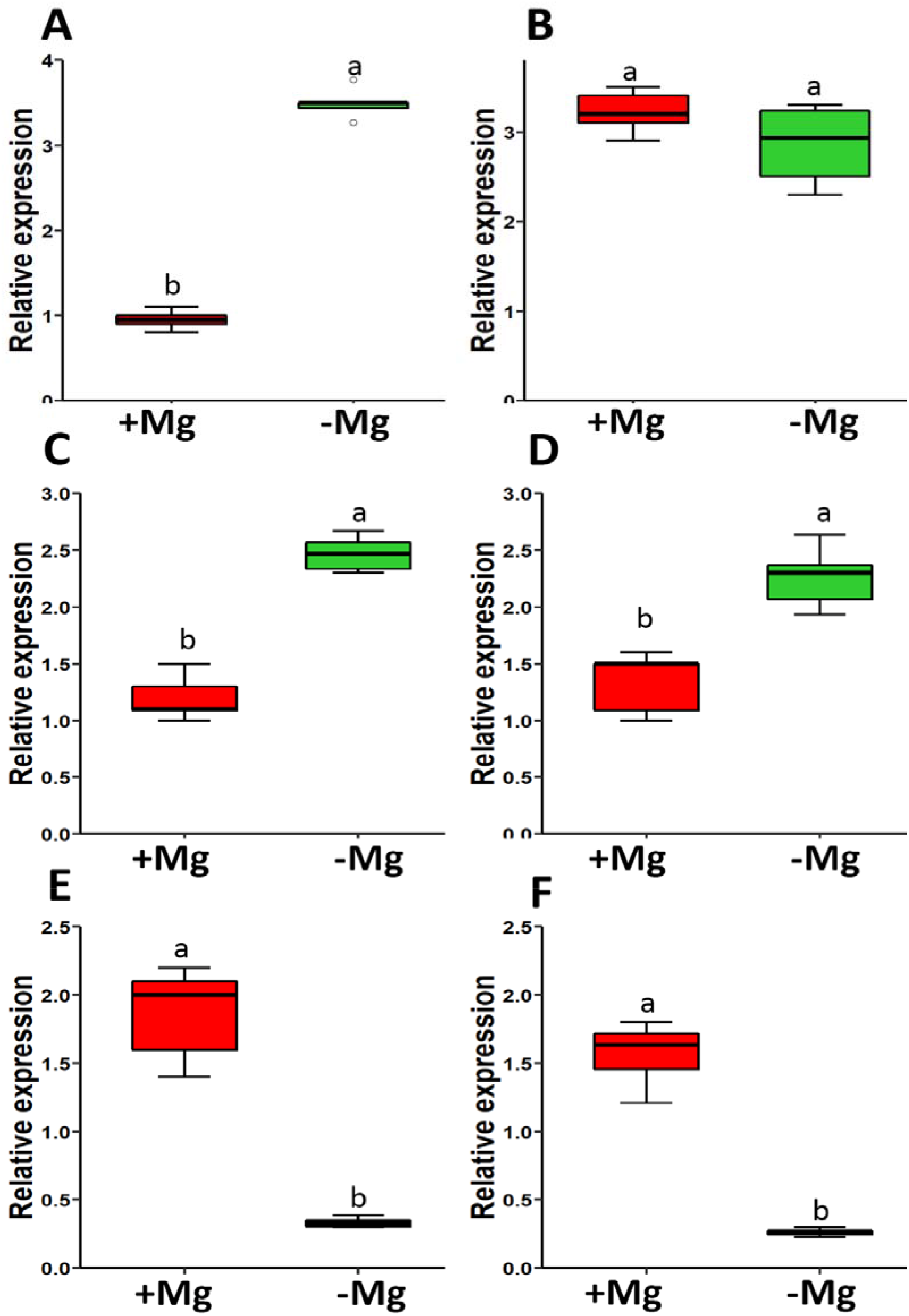
Expression level of karrikins marker genes in *Arabidopsis* grown under Mg deficient medium. (**a**) *KUF1*. (**b**) *DLK2*. (**c**) *KAI2*. (**d**) *MAX2*. (**e**) *SMAX1*. (**f**) *SMAX2*. Three-week-old seedlings were transferred to a medium have Mg (+Mg) or without Mg (-Mg) for 12 hours. The root samples were used for RNA extraction. *AtACT2* was used as a reference gene in RT-qPCR, and data gene expression was analyzed by the 2^−ΔΔCt^ method. Kruskal-Wallis’s test and post hoc Student’s t-test (P<0.05) were performed to find the significant difference between different groups; different alphabets indicate different statistical groups.

Ethylene plays a vital role in regulating RH elongation in nutrient deficient or sufficient conditions (Liam, 2001; Miao *et al*., 2017; Miao *et al*., 2018; Ru□z□ic□ka *et al*., 2007; Tian *et al*., 2009). We found that the treatment of ethylene (100 mM ethephon) recovered the root hairs in *kai2* and *max2* under Mg deficient medium (Fig. **3a, b**). Furthermore, the exogenous application of AgNO_3_ (ethylene biosynthesis inhibitor) hindered the root hair elongation of the *smax1* and *smxl2 Arabidopsis* (Fig. S1a, b). We also determined the ethylene level in WT, *kai2, smax1*, and *smxl2 Arabidopsis* grown in Mg deficient medium. The results revealed that *kai2* and *max2* had lower, while *smax1* and *smxl2* had a higher ethylene level than WT under Mg deficiency (Fig. **3c**). Ethylene biosynthesis requires ACC-synthase (ACS), and ACC oxidase (ACO) proteins (Cui *et al*., 2018; Philippos and John, 1991; Thomas *et al*., 1979), mutations in these genes are reported to have a critical role in RHL in *Arabidopsis* (Cui *et al*., 2018; Liam, 2001; Miao *et al*., 2018). A recent report showed that karrikins target genes *SMAX1* and *SMXL2* might require *ACS7* to regulate root hairs elongation in *Arabidopsis* (Carbonnel *et al*., 2020), and *acs7* mutants were unable to elongate RH in Mg deficient medium (Miao *et al*., 2018). Thus, we tested whether the exogenous application of KAR_1_ can recover RH in *acs7* or *etr1 Arabidopsis* under Mg deficient conditions. Results showed that KAR_1_ could not recover RH in *acs7 or etr1 Arabidopsis* under Mg deficient conditions (Fig. **3d, e**). We determined the relative expression level of ethylene biosynthesis genes, such as *ACS7* and *ACO1*, and ethylene signalling gene such as *ETR1*. Results revealed that *ACS7, ACO1* and *ETR1* were significantly decreased in *kai2* or *max2* while induced in *smax1* and *smxl2 Arabidopsis* compared to WT grown in low Mg conditions (Fig. **3f-h**). Overexpression of *KAI2* and *MAX2* promoted the relative expression levels of *ACS7, ACO1*, and *ETR1*, which was significantly low in *SMAX1* and *SMXL2* overexpressed *Arabidopsis* (Fig. S2). Additionally, we tested the effect of KAR_1_ application on the expression level of *ACS7, ACO1*, and *ETR1* in *Arabidopsis* grown in an Mg deficient medium. Results revealed that KAR1 promoted the expression level of *ACS7* and *ACO1* and *ETR1* under Mg-deficient conditions (Fig. S3), suggesting that *KAI2-KL* signalling promoted the root hairs elongation by increasing the ethylene biosynthesis and signalling in *Arabidopsis* under Mg deficient conditions.

**Fig. 3:**
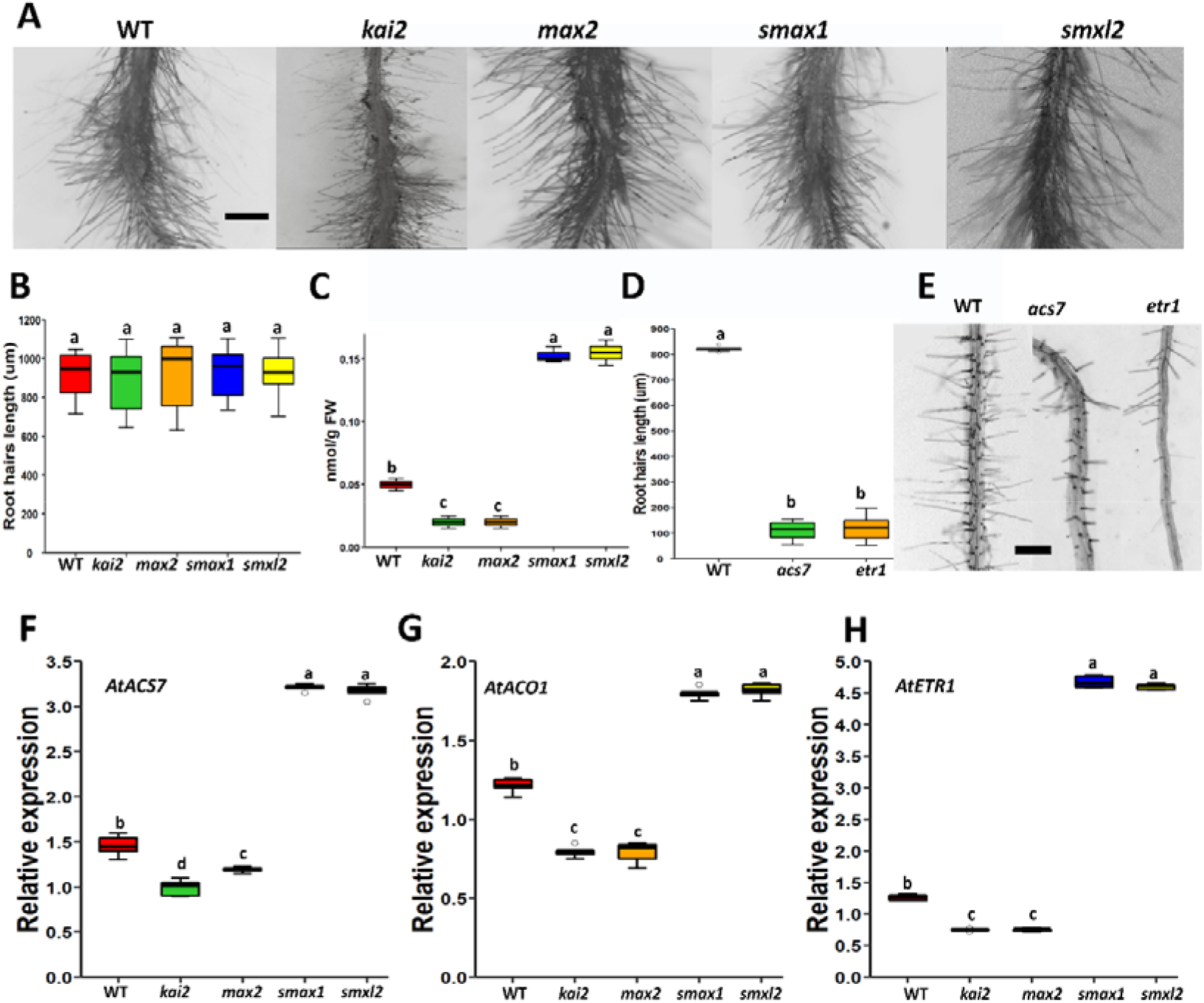
Ethylene and karrikins crosstalk in the regulation of RHL in *Arabidopsis* grow under Mg deficient conditions. (**a-b**) Ethylene effect on root hair length of karrikins signalling mutants under Mg deficient medium. (**c-d**) the effect of KAR_1_ on ethylene biosynthesis (*ACS7*) and carrier (*ETR1*) mutants *Arabidopsis* grown in Mg deficient conditions. For four days, old seedlings were shifted to -Mg media, and RHL was measured after seven days. Results are representative of 3 independent experiments. Bar, 200 μm. (**e**) ethylene level in karrikins signalling mutants under Mg deficient medium. (**f**) *ACS7*. (**g**) *ACO1*. (**h**) *ETR1*. Three-week-old seedlings were transferred to +Mg (control) or -Mg media for 12 hours. The root samples were used for RNA extraction. *AtACT2* was used as a reference gene in RT-qPCR, and data gene expression was analyzed by the 2^−ΔΔCt^ method. Kruskal-Wallis’s test and post hoc Student’s t-test (P<0.05) were performed to find the significant difference between different groups; different alphabets indicate different statistical groups. (**b, d**) n=10. (**e-h**) n=5.

Nitric oxide was another key regulator (Cristina *et al*., 2006), working simultaneously with ethylene in controlling the RHL (Miao *et al*., 2018). Therefore, we used a pharmacological technique to test whether the NO treatment could recover the root hair length of *kai2* under Mg deficiency? And whether blockers of nitric oxide synthesis could hinder the RHL of *smax1* and *smxl2* under Mg deficiency? The results showed that exogenous application of SNP (NO-releasing chemical) (Tuteja *et al*., 2004) recovered the root hairs of *kai2* under an Mg deficient medium (Fig. **4a, b**). In contrast, the exogenous application of cPTIO (NO biosynthesis inhibitor) hindered the root hair elongation of the karrikins signalling mutant *Arabidopsis* (Fig. S4). Furthermore, exogenous application of KAR1 was unable to recover RH elongation in mutants of NO biosynthesis *nos1* and carrier *nia1nia2* under Mg deficiency (Fig. **4c, d**). Moreover, we tested NO’s level in karrikins signalling mutants grown in Mg starvation conditions. Results showed that *kai2* and *max2* accumulated less. In contrast, *smax1* and *smxl2* accumulated higher NO levels than WT (Fig. **4e**), suggesting that *KAI2-KL* signalling also required NO biosynthesis in the process of RH elongation in *Arabidopsis* in response to low Mg level.

**Fig. 4:**
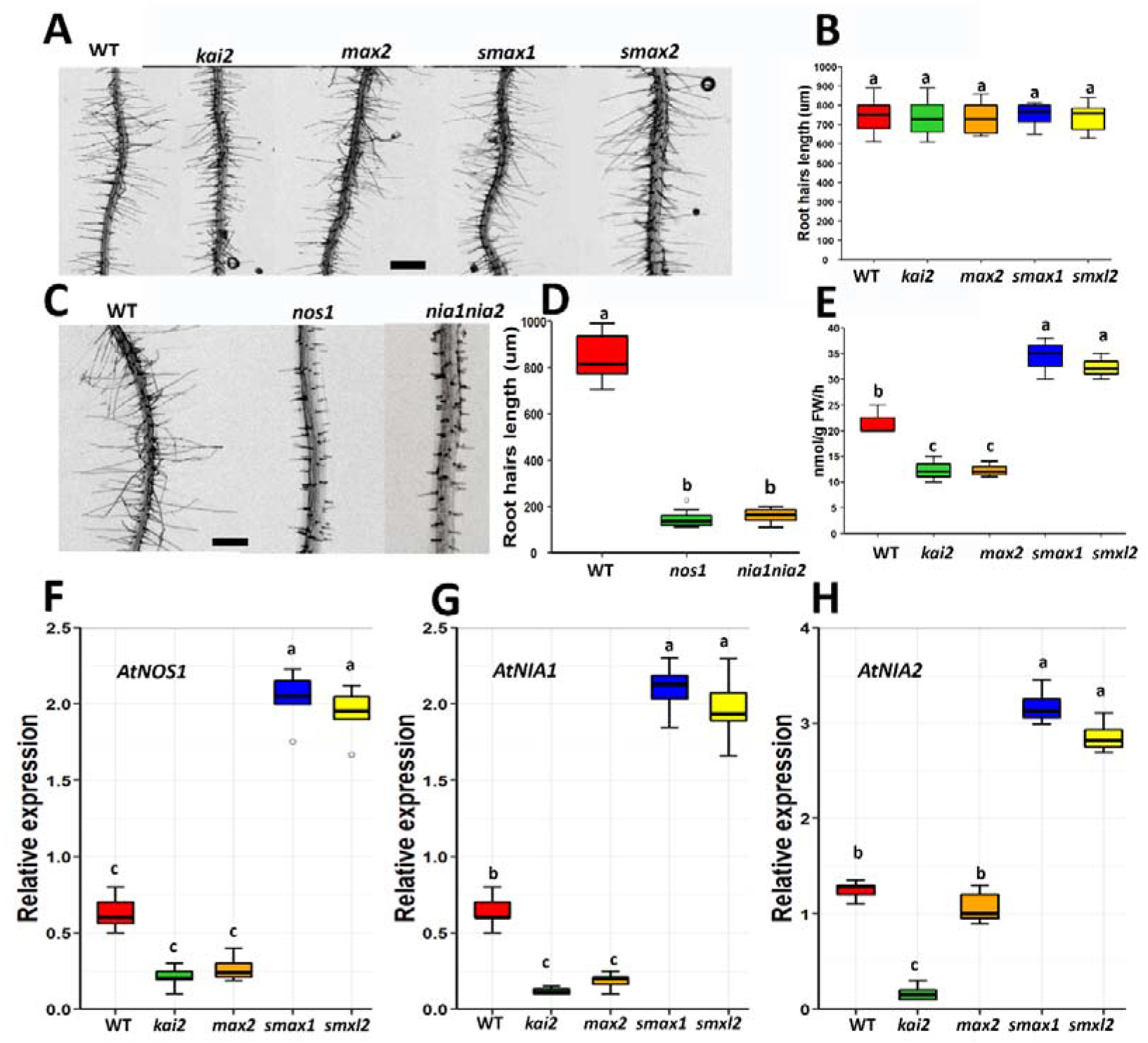
Nitric oxide and karrikins crosstalk in the regulation of RHL in *Arabidopsis* grow under Mg deficient conditions. (**a-b**) NO effect on root hair length of karrikins signalling mutants under Mg deficient medium. (**c-d**) Effect of KAR_1_ on NO biosynthesis genes mutants such as *nos1* and *nia1nia2 Arabidopsis* grown in Mg deficient conditions. For four days, old seedlings were grown in either +Mg (control) or -Mg media for seven days. Results are representative of 3 independent experiments. Bar, 200 μm. (**e**) NO level in karrikins signalling mutants under Mg deficient medium. (**f**) *NOS1*. (**g**) *NIA1*. (**h**) *NIA2*. Three-week-old seedlings were transferred to +Mg (control) or -Mg media for 12 hours. The root samples were used for RNA extraction. *AtACT2* was used as a reference gene in RT-qPCR, and data gene expression was analyzed by the 2^−ΔΔCt^ method. Kruskal-Wallis’s test and post hoc Student’s t-test (P<0.05) were performed to find the significant difference between different groups; different alphabets indicate different statistical groups. (**b, d**) n=10. (**e-h**) n=5.

NO biosynthesis gene *NOS1* has been reported to mediate the RH elongation in *Arabidopsis* under Mg deficiency (Miao *et al*., 2018). To test whether karrikins signalling RH elongation under low Mg availability is mediated by NO *biosynthesis*, we analyzed the relative expression level of NO biosynthesis genes such as *NOS1, NIA1*, and *NIA2* in karrikins signalling mutants *Arabidopsis* grown in Mg deficient medium. Results showed that the transcript level of *NOS1, NIA1*, and *NIA2* was not altered in *kai2 or max2* mutants but significantly induced in *smax1* and *smxl2* mutants under Mg deficiency (Fig. **4f-h**). Overexpression of *KAI2* and *MAX2* promoted the expression level of *NOS1, NIA1*, and *NIA2*, while overexpression of *SMAX1* and *SMXL2* reduced the relative expression level of *NOS1, NIA1*, and *NIA2* (Fig. S5). Furthermore, we tested the effect of KAR1 treatment on the expression level of *NOS1, NIA1, and NIA2* in *Arabidopsis* grown in an Mg deficient medium supplemented. The results showed that KAR_1_ promoted the *NOS1* transcript accumulation in *Arabidopsis* growing under low levels of Mg (Fig. S 6), suggesting that the karrikins signalling pathway might regulate RHL under low Mg availability through induction in NO biosynthesis in the roots of *Arabidopsis*.

Auxin is a crucial hormone that regulates RHL, and it acts downstream of NO and ethylene in RHL in response to low Mg availability (Miao *et al*., 2018). We used pharmacological technique to test whether exogenous supplementation of auxin could recover the root hairs length of *kai2*, and auxin transport inhibitor “TIBA” (Hay, 1956) could hinder the root hairs length of *smax1* and *smxl2* under Mg deficiency. Results showed that the IAA treatment recovered the root hairs of *kai2 and max2 Arabidopsis* grown in an Mg deficient medium (Fig. **5a, b**). In contrast, the exogenous application of TIBA hindered the root hair elongation of the *smax1* and *smxl2 Arabidopsis* grown in an Mg deficient medium (Fig. S7). Moreover, exogenous application of KAR_1_ was unable to promote RH elongation in *Arabidopsis* mutant in *yuc3, aux1*, and *pin2* grown in a low Mg medium (Fig. **5c, d**). Moreover, we determined the auxin level in WT, *kai2, smax1*, and *smxl2;* results showed that *kai2* had a lower, while *smax1* and *smxl2* had a higher level of auxin as compared with WT under Mg deficiency (Fig. **5e**). The relative expression level of *YUC3* was significantly reduced in *kai2* and *max2*, while *smax1* and *smxl2* had significantly higher relative expression levels of *YUC3* as compared with WT (Fig. **5f**). Moreover, the relative expression level of auxin biosynthesis gene *YUC3* was reduced in *kai2* and *max2* mutants *Arabidopsis* under Mg deficiency, which was significantly induced in *smax1and smxl2* mutants *Arabidopsis* (Fig. **5g**). Furthermore, the transcript level of the auxin efflux carrier *PIN2* was reduced in *kai2* and *max2* mutants under Mg deficiency, which was significantly induced in *smax1* and *smxl2* mutants (Fig. **5h**). Overexpression of *KAI2* and *MAX2* promoted the expression level of *AUX1, PIN2*, and *YUC3*, while overexpression of *SMAX1* and *SMXL2* reduced the relative expression level of *YUC3, AUX1*, and *PIN2* (Fig. S8). Exogenous treatment of KAR1 promoted the relative expression level of *YUC3, PIN2*, and *AUX1* under Mg-deficient conditions (Fig. S9), suggesting the karrikins signalling pathway might regulate RHL in *Arabidopsis* in response to low Mg by modulating the auxin biosynthesis and signalling.

**Fig. 5:**
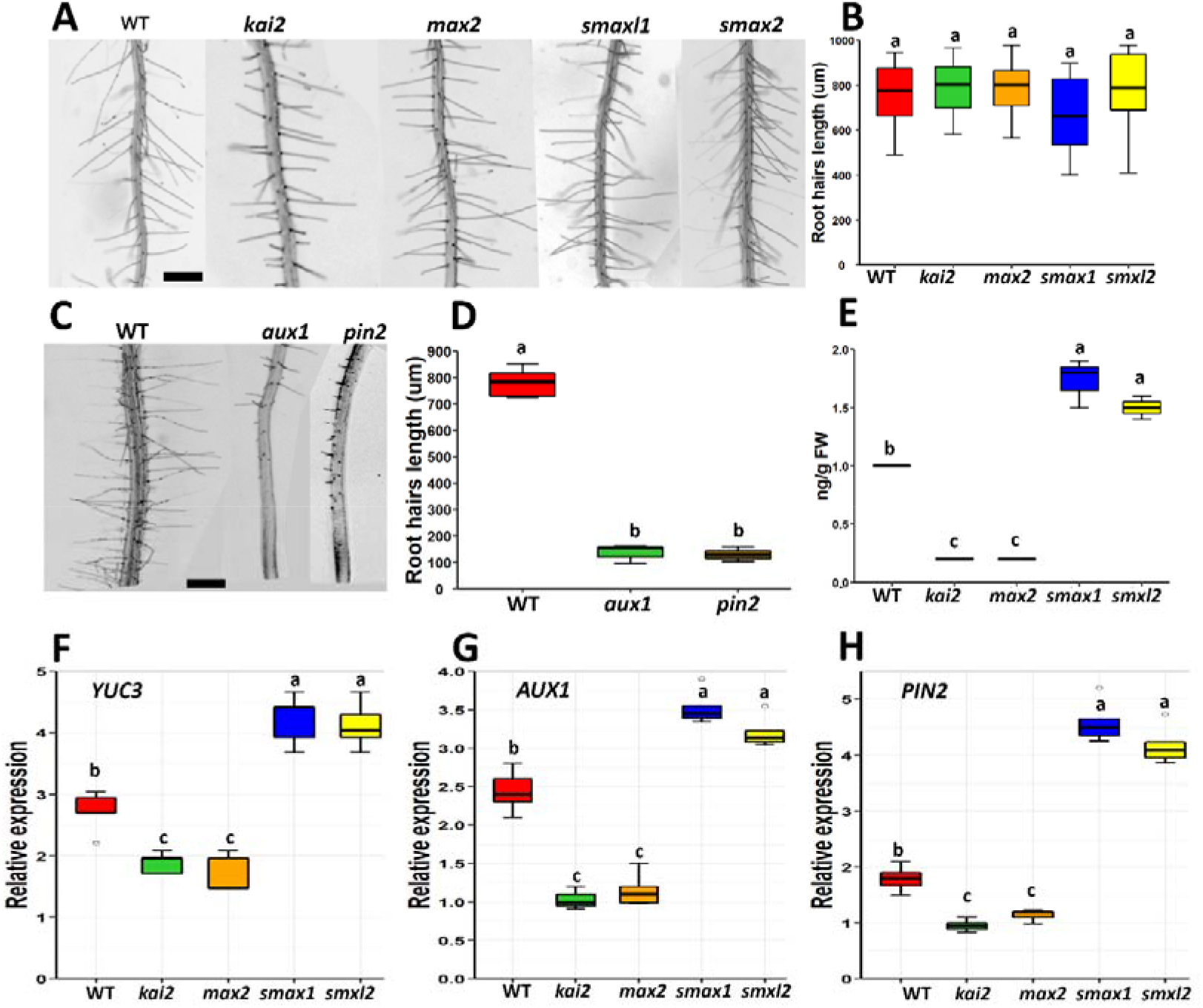
Auxin and karrikins crosstalk in the regulation of RHL in *Arabidopsis* grow under Mg deficient conditions. (**a-b**) Auxin effect on root hair length of karrikins signalling mutants under Mg deficient medium. (**c-d**) the effect of KAR_1_ on RH elongation in genes *aux1* and *pin2* mutants *Arabidopsis* grown in Mg deficient conditions. For four days, old seedlings were grown in either +Mg (control) or -Mg media for seven days. Results are representative of 3 independent experiments. Bar, 200 μm. (**e**) IAA level in karrikins signalling mutants under Mg deficient medium. (**f**) *YUC3*. (**g**) *AUX1*. (**h**) *PIN2*. Three-week-old seedlings were transferred to +Mg (control) or -Mg media for 12 hours. The root samples were used for RNA extraction. *AtACT2* was used as a reference gene in RT-qPCR, and data gene expression was analyzed by the 2^−ΔΔCt^ method. Kruskal-Wallis’s test and post hoc Student’s t-test (P<0.05) were performed to find the significant difference between different groups; different alphabets indicate different statistical groups. (**b, d**) n=10. (**e-h**) n=5.

## Discussion

Root hair elongation (RHL) is a crucial adaptation which provides more surface area for nutrient absorption to plants facing nutrient deficiency. Under Mg deficiency, RHL is coordinately regulated by hormones such as auxin, ethylene, and nitric oxide (Miao *et al*., 2018; Niu *et al*., 2014). This study found that the *KAI2-KL* signalling involved root hair elongation in response to Mg starvation. Moreover, this study revealed the involvement of auxin, ethylene, and nitric oxide in karrikins regulation of RHL under Mg deficiency.

Magnesium (Mg) is a secondary nutrient required for plant growth and development, and Mg starvation could cause yield reduction in crops (Christian *et al*., 2010). Therefore, Mg starvation is often considered a severe limitation in crop production (Petra and Zed, 2012; Xie *et al*., 2021). Elongation of RH is an adaptive mechanism in plants for survival in low Mg availability (Niu *et al*., 2014). Plant hormones are essential for RHL under low Mg availability (Mao *et al*., 2014; Miao *et al*., 2018; Niu *et al*., 2014). Our results showed that the positive regulators of karrikins *kai2* and *max2* mutant *Arabidopsis* was deficient in root hair elongation under Mg deficiency. While mutation in *smax1* and *smxl2*, which are negative regulators of karrikins, had long root hairs under Mg deficiency (Fig. **1a-d**). These results were a hint showing the involvement of *KAI2-KL* signalling in the regulation RH elongation in response to Mg deficiency, which agreed with the previous results showing that *KAI2-Kl* signalling regulated the RH elongation under control or a macro-element such as phosphorous deficiency (Carbonnel *et al*., 2020; José Antonio *et al*., 2022). Interestingly, exogenous treatment of pharmacological karrikins, such as KAR_1_ or KAR_2_, induced the root hairs elongation in *Arabidopsis* growing in Mg starvation conditions (Fig. **1e-f**). These results verified that KL or its synthetic alternates might induce RHL in *Arabidopsis* grown in Mg deficient conditions, which was in agreement with previous reports showing that the plant activates the RH regulatory genes as a molecular response to a nutrients deficiency (José Antonio *et al*., 2022; Kumar *et al*., 2015; Miao *et al*., 2018; Michael *et al*., 2009). Furthermore, Mg deficiency significantly promoted the relative expression level of *KAI2* and *MAX2*. In contrast, *SMAX1* and *SMXL2* expression levels were reduced in Mg deficient medium compared with Mg sufficient medium (Fig. **2**), suggesting that *KAI2-KL* signalling is involved in RH elongation under Mg deficiency.

Ethylene has a significant role in the regulation of RHL under control or nutrients deficient conditions (Carbonnel *et al*., 2020; Jason *et al*., 1998; Miao *et al*., 2018; Mimi *et al*., 1995; Ru□z□ic□ka *et al*., 2007; Tian *et al*., 2009; Zhu *et al*., 2006). We found that Ethephon (ethylene releasing compound) treatment recovered the root hairs of *kai2* under Mg deficient medium (Fig. **3a, b**). In contrast, the exogenous application of AgNO_3_ (ethylene biosynthesis inhibitor) hindered the root hair elongation in *smax1* and *smxl2 Arabidopsis* (Fig. S1a, b), suggesting a blatant interference of ethylene in *KAI2-Kl* signalling induced RHL in *Arabidopsis* grown in low Mg conditions. Furthermore, previous reports showed that Mg deficiency stimulated ethylene production via activating the ACS proteins, essential for ethylene biosynthesis (Chae and Kieber, 2005; Yamagami *et al*., 2003). previous reports showed that the relative expression level of the ACS family members, such as *ACS7*, and ethylene oxidase genes, such as *ACO1*, was induced in the roots of *Arabidopsis* growing in an Mg deficient medium (Carbonnel *et al*., 2020; Miao *et al*., 2017; Miao *et al*., 2018). The results of our study agreed with the reports mentioned above, showing that KAR_1_ treatment could not recover the RHL in *acs7, aco1*, or *etr1* mutants (Fig. **3c, d**). Furthermore, overexpression of *AtKAI2* and *AtMAX2* in *Arabidopsis* promoted the expression level of *ACS7, ACO1or ETR1* in response to Mg deficiency. In contrast, overexpression of *AtSMAX1* and *AtSMXL2* in *Arabidopsis* suppressed the expression level of *ACS7, ACO1or ETR1* in response to Mg deficiency (Fig. S2), suggesting that KL signalling regulates ethylene signalling in *Arabidopsis* in response to Mg deficiency. Exogenous application of KAR_1_ also strengthened the above results and induced the expression level of *ACS7, ACO1or ETR1* in *Arabidopsis* grown in low Mg conditions (Fig. S3). Previous studies showed that *acs7, aco1*, or *etr1* are defective in ethylene biosynthesis or signalling, so these have short RHL under Mg stress or control conditions (Liam, 2001; Miao *et al*., 2017; Mimi *et al*., 1995). Furthermore, reduced ethylene level in *kai2* and *max2* mutants as compared to WT and an induced ethylene level in *smax1* and *smxl2* in Mg deficient medium (Fig. **3e**), showing the need for ethylene in *KAI2-Kl* signalling regulation of RHL in *Arabidopsis* under Mg deficient conditions as reported in previous studies (Carbonnel *et al*., 2020; Miao *et al*., 2018). Previous reports showed that *ACS7, ACO1*, and *ETR1* are the critical genes in ethylene synthesis and signalling and regulate RHL in *Arabidopsis* grown in Mg starvation (Miao *et al*., 2017; Miao *et al*., 2018). The promoted relative expression of *ACS7, ACO1, and ETR1* in *smax1*and *smxl2* mutants (Fig. **3f-h**), was agreed with previous studies (Miao *et al*., 2017; Miao *et al*., 2018) and could be a reason for elongated RH in *smax1* or *smaxl2 Arabidopsis* grown under Mg starvation. These results suggest that *KAI2-KL* signalling mediates ethylene biosynthesis resulting in the induction of RHL in *Arabidopsis* (Carbonnel *et al*., 2020) grown under Mg stresses (Miao *et al*., 2017).

Nitric oxide works simultaneously with ethylene in the process of root hair elongation (Cristina *et al*., 2006; Miao *et al*., 2018). Under Mg deficient conditions, plants have developed an adoptive mechanism to lessen cellular damage (Christian *et al*., 2010). The previous report showed that NO-mediated the RH elongation under Mg deficient conditions, pharmacological treatments of NO inhibitor cPTIO, and the mutation in NO regulatory genes significantly reduced the RH elongation under Mg deficient conditions (Miao *et al*., 2018). Furthermore, exogenous supplementation of NO donor SNP has similar RHL to the plants grown in Mg-deficient conditions; and the NO insensitive mutants were unable to RH elongate under Mg deficient conditions (Miao *et al*., 2018). This study revealed that exogenous application of SNP could recover the RHL of *kai2* under Mg deficiency, suggesting that *kai2* might not contain enough NO level to elongate root hairs under Mg deficiency. At the same time, a mutation in *smax1* and *smxl2* had higher level NO levels and produced long root hairs in response to Mg starvation (Fig. **4**). These results agreed with the previous reports showing that exogenous application of SNP, a NO-releasing chemical (Tuteja *et al*., 2004), recovered RHL in *Arabidopsis* in an Mg deficient medium (Miao *et al*., 2018). Furthermore, *kai2* and *max2* had a lower level of NO, which agreed with the previous studying showing that karrikins signalling genes were reported to have a low level of NO. Exogenous application of cPTIO blocked the root hair elongation in *smax1 and smxl2* (Fig. S4), suggesting NO’s importance in KL signalling induced RHL in response to low Mg. Another report showed that KAR1 caused NO production in *S. miltiorrhiza* hairy root (Zhou *et al*., 2019). We found that overexpression of *AtKAI2* and *AtMAX2* mediated RHL in *Arabidopsis* in response to low Mg levels by suppressing NO biosynthesis by decreasing the expression level of *NOS1, NIA1*, and *NIA2* (Fig. S5). Moreover, exogenous supplementation of KAR_1_ could not promote the RHL in *nos1* and *nia1nia2* double mutant *Arabidopsis* grown in Mg deficient conditions (Fig. **4c, d**). Additionally, the transcript level of NO biosynthesis genes such as *NOS1, NIA1*, and *NIA2* was not changed in *kai2* mutants under Mg deficiency, which was significantly induced in *smax1*and *smxl2* mutants (Fig. **4f-h**). KAR_1_ treatment induced the transcript level of *NOS1, NIA1*, and *NIA2* under Mg-deficient conditions (Fig. S6), which might provide the evidence that karrikins might regulate root hairs length by modulating NO biosynthesis or signalling.

Auxin is also reported as a downstream player of ethylene and NO (Miao *et al*., 2018). Previous reports showed that *KAI2* regulation of RHL is controlled by auxin signalling. Thus, we tested the role of auxin biosynthesis and signalling in *KAI2-KL* signalling regulation of RHL in *Arabidopsis* grown under Mg deficiency. Therefore, exogenous application of auxin restored RHL in *kai2* and *max2* mutants, same as wild-type under the medium level of Mg (Fig. **5a, b**), suggesting that mutation in *kai2* and *max2* lead to a shortage in indigenous auxin or auxin signalling. *YUCCA3* (*YUC3*) regulates auxin biosynthesis in roots (Cao *et al*., 2019; Chen *et al*., 2014). AUX1 is an auxin export carrier protein that metabolizes 2,4-dichlorophenoxyacetic acid (2,4-D), while PIN-family proteins use auxin analogue 1-naphthalene acetic acid (NAA) as its substrate (Ma *et al*., 2017). Exogenous application of KAR1 did not recover the reduced RHL in *yuc3, aux1*, or *pin2 Arabidopsis* (Fig. **5c, d**), suggesting the pivotal role of auxin in *KAI2-Kl* signalling regulation of RHL in Mg deficient conditions. A reduced level of IAA in *kai2* and *max2 Arabidopsis* (Fig. **5e**) suggests that the short length of *kai2* and *max2* RH might be caused by reduced auxin level in the roots of the *Arabidopsis* grown at a low Mg level (Bhosale *et al*., 2018). The expression level of *YUC3, PIN2*, and *AUX1* in *kai2* and *max2* was not changed significantly as compared with wild-type *Arabidopsis* (Fig. **5f-h**), which suggests that the karrikins signalling pathway regulates the auxin biosynthesis or transport, these results agreed with the previous reports (Chen *et al*., 2014; José Antonio *et al*., 2022).

Ethylene facilitated NO biosynthesis by activating *NOS1* in the roots of *Arabidopsis* grown in an Mg-deficient medium. Moreover, NO boosted ethylene production by promoting the expression level of ACC synthases (ACS) in *Arabidopsis* roots grown in an Mg-deficient medium (Miao *et al*., 2017). Another report showed auxin promoted NO production by stimulating NO biosynthesis genes (Cristina *et al*., 2006). Ethylene induced auxin biosynthesis and signalling during regulation of RHL in Arabidopsis (Dubois *et al*., 2018; Ru□z□ic□ka *et al*., 2007). Another report also revealed a thought-provoking reciprocal regulation of ethylene and auxin synthesis; and auxin increased the transcription of genes regulating ethylene biosynthesis (Swarup *et al*., 2007). The *kai2* and *max2* have short RHL in response to Mg starvation might be due to the lower level of ethylene, NO, and IAA. In contrast, induction of RHL in *smax1* and *smxl2* might be because of a higher level of auxin, ethylene, or NO, suggesting that *KAI2-KL* signalling interacts with ethylene, NO, and auxin biosynthesis in the process of RH elongation in response to low Mg availability. Furthermore, the inability of KAR_1_ to recover RHL in *acs7, etr1, nos1, nia1nia2,pin2, yuc3*, and *aux1* grown in Mg starvation showed that *KAI2-Kl* signalling might work at upstream of ethylene, auxin, or NO signalling during the process of RH elongation in response to low Mg availability.

Conclusively, this study revealed the molecular mechanism of *KAI2-KL* signalling in RH elongation in response to Mg deficiency. Mg starvation transcriptionally activated the *KAI2* and *MAX2* through previously reported Mg signalling pathways (Miao *et al*., 2018), probably enhancing the sensitivity to KL. The activated KL signalling pathway could increase the degradation of *SMAX1* and *SMXL2*, resulting in the stimulated expression levels of *ACS7, ACO1, and ETR1* (Carbonnel *et al*., 2020), thus promoting ethylene biosynthesis in the root of *Arabidopsis* grown in an Mg deficient medium. Induced levels of ethylene might cause an induction in NO and auxin accumulation in the roots under low Mg conditions. Consistent with this model, RHL was promoted by KAR_1_, KAR_2_, ethephon, SNP, and NAA in *Arabidopsis* grown in Mg deficient conditions (Fig. **6**). Studying whether this regulatory network physically interacts with the *KAI2-KL* signalling pathway will be thought-provoking. Moreover, if KL were found in the future, it will be exciting to examine whether Mg availability regulates KL biosynthesis.

**Fig. 6:**
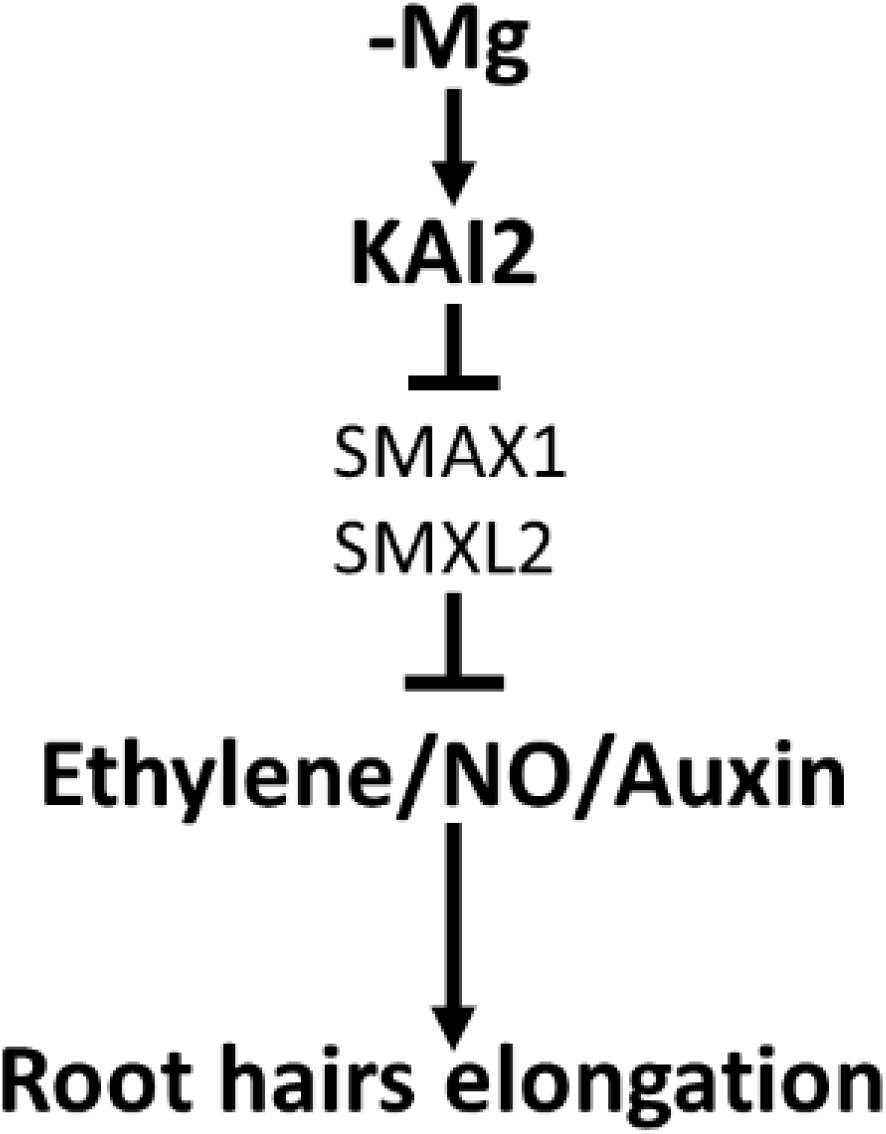
Model representing the mechanism in *KAI2* regulation of RHL in response to Mg deficiency. The model displayed that Mg deficiency promoted *KAI2* signalling, and inducement in *KAI2* led to downregulation of *SMAX1* and *SMXL2*. Ethylene, NO, and auxin biosynthesis and signalling were induced when *SMAX1* and *SMXL2* were present in low concentrations. Induction in ethylene, NO, and auxin biosynthesis and signalling promoted root hair elongation.

## Supporting data

**Table S1**: List of primers

**Fig. S1:** Effect of AgNO_3_ on the root hair elongation in KL-signaling mutants *Arabidopsis* in Mg starvation conditions

**Fig. S2:** Overexpression of karrikins signaling genes in Arabidopsis affected the expression level of *ACS7, ACO1*, and *ETR1* in *Arabidopsis* grown in low Mg conditions.

**Fig. S3:** Effect of KAR_1_ application on the expression level of *ACS7, ACO1*, and *ETR1* in *Arabidopsis* grown in low Mg conditions.

**Fig. S4:** Effect of cPTIO on root hair length of karrikins signaling mutants under Mg deficient condition.

**Fig. S5:** Overexpression of karrikins signaling genes in *Arabidopsis* affected the expression level of *NOS1, NIA1*, and *NIA2* in *Arabidopsis* grown in low Mg conditions.

**Fig. S6:** Effect of KAR_1_ application on the expression level of *NOS1, NIA1*, and *NIA2* in *Arabidopsis* grown in low Mg conditions.

**Fig. S7:** Effect of TIBA on the root hairs length in KL-signaling mutants *Arabidopsis* in Mg starvation conditions

**Fig. S8:** Overexpression of karrikins signaling genes in Arabidopsis affected the expression level of *YUC3, AUX1*, and *PIN2* in *Arabidopsis* grown in low Mg conditions.

**Fig. S9:** Effect of KAR_1_ application on the expression level of *YUC3, AUX1*, and *PIN2* in *Arabidopsis* grown in low Mg conditions.

## Data availability

All data are available within the paper, and its supplementary materials are published online.

## ACKNOWLEDGEMENTS

This work was funded by the Anhui Natural Science Foundation funded this study (1908085QC141&1808085QC73), the National Key R&D Program of China (2018YFD0502001&2018YFD0300901), the Science and Technology Service program of the Chinese Academy of Sciences (KFJ-STS-ZDTP-054), the key program of the 13th five-year plan, CASHIPS (No. KP-2019-21), the STS supporting project of the Chinese Academy of Sciences in Fujian Province (2019T3031), the project of the Dean Fund of Hefei Institute of Physical Science, Chinese Academy of Sciences, (YZJJ2020QN29), general Project of Natural Science Foundation of Anhui Province (2008085MC60), major science and technology project of Anhui Province (202103a06020002), the Chinese Academy of Sciences Plant Resources Innovation Platform Phase II (KFJ-BRP-007-014), the Anhui Province Directed Entrustment Project (2021d06050003).

## Contributions

FAS and JN designed the experiment. And FAS, XC, and CGT performed the experiments in the whole study. FAS did microscopy, arranged the data, and performed the statistical analysis. FAS and JN drafted the manuscript; all authors critically reviewed the final manuscript.

